# The Carcinogenome Project: In-vitro Gene Expression Profiling of Chemical Perturbations to Predict Long-Term Carcinogenicity

**DOI:** 10.1101/323964

**Authors:** Amy Li, Xiaodong Lu, Ted Natoli, Joshua Bittker, Nisha Sipes, Aravind Subramanian, Scott Auerbach, David H. Sherr, Stefano Monti

**Affiliations:** Computational Biomedicine, Boston University School of Medicine, Boston, MA, USA; Cancer Program, Broad Institute of MIT and Harvard, Cambridge, MA, USA; Toxicoinformatics Group, National Institute of Environmental Health Sciences, Durham, NC, USA; Department of Environmental Health, Boston University School of Public Health, Boston, MA, USA; Bioinformatics Program, Boston University, Boston, MA, USA

## Abstract

**Background:** Most chemicals in commerce have not been evaluated for their carcinogenic potential. The current de-facto gold-standard approach to carcinogen testing adopts the two-year rodent bioassay, a time consuming and costly procedure. Alternative approaches, such as high-throughput in-vitro assays, show promise in addressing the limitations in carcinogen screening.

**Objectives:** We developed a screening process for predicting chemical carcinogenicity and genotoxicity and characterizing modes of actions (MoAs) using in-vitro gene expression assays.

**Methods:** We generated a large toxicogenomics resource comprising ~6,000 expression profiles corresponding to 330 chemicals profiled in HepG2 cells at multiple doses and in replicates. Predictive models of carcinogenicity were built using a Random Forest classifier. Differential pathway enrichment analysis was performed to identify pathways associated with carcinogen exposure. Signatures of carcinogenicity and genotoxicity were compared with external data sources including Drugmatrix and the Connectivity Map.

**Results:** Among profiles with sufficient bioactivity, our classifiers achieved 72.2% AUC for predicting carcinogenicity and 82.3% AUC for predicting genotoxicity. Our analysis showed that chemical bioactivity, as measured by the strength and reproducibility of the transcriptional response, is not significantly associated with long-term carcinogenicity, as evidenced by the many carcinogenic chemicals that did not elicit substantial changes in gene expression at doses up to 40 μM. However, sufficiently high transcriptional bioactivity is necessary for a chemical to be used for prediction of carcinogenicity. Pathway enrichment analysis revealed several pathways consistent with literature review of pathways that drive cancer, including DNA damage and DNA repair. These data are available for download via https://clue.io/CRCGN_ABC, and a web portal for interactive query and visualization of the data and results is accessible at https://carcinogenome.org.

**Conclusions:** We demonstrated a short-term in-vitro screening approach using gene expression profiling to predict long-term carcinogenicity and infer MoAs of chemical perturbations.

## Introduction

Despite significant investments into cancer research over the last decades, approximately 1.7 million new cancer cases and 600k cancers deaths were estimated in the U.S. in 2017 alone (American Cancer Society 2017). Of these, 90-95% are not attributable to known heritable genetic factors, thus making environmental exposures a major suspect in driving cancer risk (Anand et al. 2008), notwithstanding recent studies pointing to the rate of cell replications as an important determinant of cancer risk variation among different tissue types (Tomasetti and Vogelstein 2015; Tomasetti et al. 2017). Most research aimed at assessing cancer risk from exposure has primarily relied on epidemiological studies of past human exposures to suspected carcinogens in cancer clusters, and on carcinogen screening based on the 2-year rodent-based bioassay. Epidemiological studies rely on observational data, and as such it is often difficult to rule out the possibility of spurious associations due to confounding effects. They also require that exposure to a suspected carcinogen is documentable. Even when the nature of the chemical exposure and the exposure dose is known, epidemiological studies require long follow-up periods, hence are not appropriate for the evaluation of new chemicals on the market. Similarly, the 2-year rodent bioassay, the gold standard for carcinogen testing, is time-consuming and requires up to $4 million and more than 800 animals per compound. As a result, less than 2% of the ~85,000 chemicals registered in the TSCA Chemical Substance Inventory have been tested by this approach (Bucher and Portier 2004; Gold et al. 2005; Huff et al. 2008).

High-throughput transcriptional profiles from short-term chemical exposures have proven useful for predicting long-term carcinogenicity and for capturing multiple biological MoAs of long-term carcinogenicity. Many studies have explored the use of high-throughput transcriptional profiling in rodent models (Eichner et al. 2013; Ellinger-Ziegelbauer et al. 2008; Gusenleitner et al. 2014; Kossler et al. 2015; Uehara et al. 2011). However, questions remain about the relevance of rodent models for characterizing human carcinogenicity, and most importantly, they are still excessively time-consuming and expensive for large-scale testing. In-vitro-based screens would help address the time and cost constraints of carcinogen testing through automated high-throughput plating, exposure treatment, and assaying, and would address the human relevance concern by relying on human cell lines that match the biological contexts of human populations at risk. EPA’s Toxcast (Judson et al. 2010; Richard et al. 2016) and Tox21 initiatives (Schmidt 2009; Tice et al. 2013) have used various reporter assays to characterize adverse effects across thousands of in-vitro chemical exposures. However, while these efforts use high-throughput techniques with carefully selected gene, pathway and adverse-response-centric endpoints, the number of assays and the diversity of endpoints are limited. For instance, ToxCast uses 624 in-vitro endpoints mapped to 315 genes in Phase I (Judson et al. 2010) and an additional ~200 new endpoints in Phase II (Richard et al. 2016). Studies utilizing this data for the assessment of chemical carcinogenicity have emphasized the need to expand the assay set to better characterize diverse MoAs of certain carcinogens (Kleinstreuer et al. 2013). mRNA profiling, by assaying the entire transcriptome, or a large portion of it, represents a promising solution to this need by providing an agnostic view of which genes and pathways are relevant to chemical-induced carcinogenesis.

Given the technological advances in gene expression profiling and the development of cost-effective sequencing platforms, opportunities arise for their use in large-scale toxicological screenings. One such solution is the Luminex-1000 (L1000) platform (Peck et al. 2008), a low-cost, high-throughput bead-based platform that measures the expression of ~1000 landmark genes and infers the remaining genes in the transcriptome by imputation. This platform was used in the creation of the Connectivity Map (CMap) (Subramanian et al. 2017), which now includes 1.3 million perturbation profiles of drugs and small molecules and has been instrumental in the discovery of small molecule MoAs. Due to its cost-effectiveness and appropriateness for large-scale perturbation screening, we adopted it for the profiling of chemical carcinogens.

We applied the L1000 platform to study the effects of chemical perturbations of previously validated rat liver carcinogens and non-carcinogens in HEPG2 cell lines. Our approach used machine-learning techniques to build predictive models of the long-term carcinogenicity of chemicals based on L1000-derived gene expression profiles of human cell lines exposed to the studied chemicals. Furthermore, we annotated the In-vitro-derived gene signatures by performing pathway enrichment of carcinogens *vs*. non-carcinogens, to identify MoAs associated with chemical induced carcinogenesis. Signatures derived from this study were also compared to external gene signatures and chemical annotations from knowledge bases such as Drugmatrix, CMap, and Tox21, to verify the consistency of results and expand the interpretation of findings. An overview of our experimental design and analysis aims is presented in Figure S1.

## Methods

### Chemical selection and annotation

In the chemical selection process, we prioritized chemicals with long-term rodent liver carcinogenicity annotation for inclusion in this experiment. Long-term carcinogenicity annotations were derived from the Carcinogenic Potency Database (CPDB) (Fitzpatrick 2008). Additional chemicals without carcinogenicity annotation were included on the basis of interest to the Superfund Research Program (environmental toxicants), presence in controversial commercial products (included for predictive purposes), and evidence of binding to the aryl hydrocarbon receptor (AhR), as the AhR is an important mediator of xenobiotics, including carcinogens. A complete list of chemicals and their annotations is provided in Table S1. For CPDB annotations, the final carcinogenicity labels denote “+” if carcinogenic in rat liver (female or male) or “−” if non-carcinogenic in both rat and mouse (in female and male) across all tested organs in the CPDB. Genotoxicity labels denote “+” if mutagenic or weakly mutagenic in the Salmonella assay, and “−” otherwise.

### Chemical procurement and data generation

Chemicals were procured from the Tox21 library of the National Toxicology Program (NTP) when available, or from Sigma-Aldrich otherwise. Compound purity and identity were confirmed by UPLC-MS (Waters, Milford, MA). Purity was measured by UV absorbance at 210 nm or by Evaporative Light Scattering (ELSD). Identity was determined on a SQ mass spectrometer by positive and/or negative electrospray ionization. HepG2 cells (liver cancer cell line) were exposed to each chemical for 24 hours in 384-well plates in 6 doses in triplicate wells per dose and chemical combination, starting from 40μM maximum dose (40mM stock diluted 1:1000) for NTP chemicals (or 20μM for chemicals procured from Sigma-Aldrich) in series of two-fold dilutions. The sole exception to the standard dosage is 2,3,7,8-Tetrachlorodibenzo-p-dioxin (TCDD), which had a starting dose of 50nM due to its extreme potency. Following 24 hours of chemical exposure, the gene expression of the HEPG2 cells was profiled using the L1000 platform, a high-throughput assay that measures the expression of ~1000 landmark genes and computationally infers the expression of non-measured transcripts (Subramanian et al. 2017).

For each perturbation and landmark gene, we computed the change in gene expression following the perturbation using a moderated z-score procedure as described in the CMap-L1000 workflow. Differential expression values were calculated as moderated z-scores for each landmark gene and each unique perturbation (chemical and dose combination) perturbation, collapsed to a single value across replicates.

### Assessing the transcriptional strength of a perturbation

We used the *transcriptional activity score* (TAS) as a summary measure of the impact of a chemical perturbation on landmark gene expression. TAS integrates *signature strength*, defined as the number of genes up-regulated or down-regulated by a particular perturbation above a given moderated z-score threshold, and *replicate correlation*, a measurement of similarity among triplicate profiles corresponding to the same perturbation (unique combination of chemical, dose, cell line, time). Formally, TAS is quantified as the geometric mean of the signature strength (SS_ngene_) and the replicate correlation (CCq75). SS_ngene_ is defined as the number of landmark genes (<u>card</u>inality) with ModZ greater than 2, wherein ModZ is defined as the 978-element vector of replicate collapsed z-scores of landmark genes, and CCq75 is the 75th quantile of the spearman correlations between replicates in landmark space.

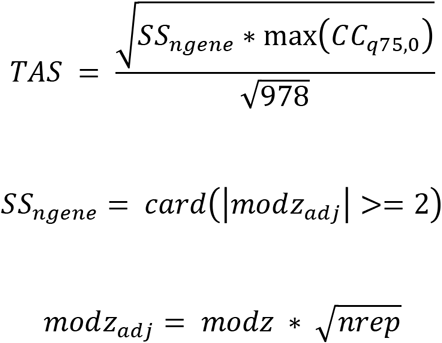

TAS is calculated for each aggregated profile (one unique score per chemical and dose combination). This metric takes value in the [0,1] range, with higher values of TAS taken to represent a higher level of chemical bioactivity.

### Statistical tests for comparison of TAS across profiles

We tested for the difference in TAS values among adjacent dose groups using a one-tailed Wilcoxon Signed-Rank Test (paired difference test), with the pairing determined by the unique chemical IDs to determine the statistical significance of strictly increased TAS levels between adjacent and increasing dose groups.

We next tested for difference in TAS between chemicals. In particular, for each dose rank, two-group comparisons of TAS scores between carcinogens and non-carcinogens, and between genotoxicants and non-genotoxicants, were conducted using one-tailed unpaired two-samples Wilcoxon test, to determine the presence and significance of increased TAS for the carcinogenic compared to non-carcinogenic group, or for the genotoxic compared to non-genotoxic group.

### Equivalent In-vitro dose (Cmax) estimation and association with TAS

We assessed the relationship between in-vitro transcriptional bioactivity (TAS) and corresponding in-vivo dose used in the rodent bioassay from which carcinogenicity labels were derived. To this end, using a toxicokinetic model (Pierce et al. 2017), we estimated the *equivalent in-vitro dose* (Cmax) corresponding to the in-vivo dose tested in the rat bioassay.

Cmax values were estimated using the R package HTTK v1.8 (Pierce et al. 2017). For carcinogenic compounds, these values were derived from the CPDB-reported median toxic dose (TD50) administered in rats. For non-carcinogenicity compounds, Cmax values were derived from the CPDB-reported maximum dose administered in rats. Chemicals with missing TD50 (if carcinogenic) or maximum dose (if non-carcinogenic) were omitted from this analysis. It is assumed that dosing was once per day for 365 days.

To determine the association between TAS, carcinogenicity, and Cmax, we used the following linear regression model:

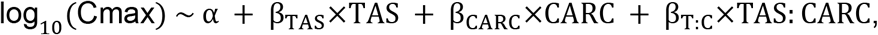

where TAS denotes the mean TAS for each chemical (across 6 doses), and CARC denotes the carcinogenicity status of the chemical in the rodent bioassay. We tested for significance of the coefficients β_TAS_, β_CARC_, β_T:C_ under the null hypotheses of zero-valued coefficients (no effect).

### Supervised learning for prediction of carcinogenicity and genotoxicity

To build classifiers for the prediction of carcinogenicity and genotoxicity, we used the moderated z-scores of landmark genes as predictive features. The Random Forest classifier was used, as implemented in the R package *caret* (Kuhn 2012). The performance of the classifier was evaluated using a resampling scheme consisting of 25 random repeats of training on 70% of the samples and testing on the remaining 30%. The training and test set split was performed at the chemical level, so that all replicates of each chemical were only included either in the train or the test set, to avoid “information leakage” (over-fitting). To assess the effect of chemicals’ bioactivity on the performance of the classifier, the evaluation was repeated on different subsets of profiles corresponding to different TAS thresholds (all profiles, TAS>0.2, >0.3, >0.4). Area under the ROC curve (AUC) was used for the assessment of a classifier performance, as it is a well-established metric that captures the trade-off between sensitivity and specificity across multiple thresholds.

Final predictions of carcinogenicity and genotoxicity were made using leave-one-(chemical)-out (LOCO) cross-validation (CV); that is, at each CV iteration, a single chemical’s profiles across multiple doses are left out and a classifier is trained based on all remaining chemicals, then applied to the prediction of the left-out chemical’s profiles. This procedure is repeated with each of the TAS subsets.

### Deriving pathway signatures of carcinogenicity

We derived pathway activity scores using the R Bioconductor package GSVA (Hänzelmann et al. 2013; Hänzelmann et al. 2014). This tool takes as input a gene-by-sample expression matrix and generates a geneset-by-sample enrichment score matrix, with its entries representing the pathway enrichment of each sample with respect to each of a user specified list of gene-sets. Pathway enrichment scores were calculated for pathways in the MsigDB C2 Reactome pathway compendium (Fabregat et al. 2017; Liberzon et al. 2011; Milacic et al. 2012). The geneset-projected matrix was then used as input for differential analysis with respect to sample phenotype labels (carcinogenicity or genotoxicity) using the R Bioconductor package *limma* (Ritchie et al. 2015; Smyth 2005) to identify pathways with differences in activity levels between chemical groups. This differential analysis was repeated from data inputs with various TAS thresholding (TAS > 0, 0.2, 0.3, 0.4). One-sided p-values consistent with the direction of change in pathway activity scores were estimated. The p-values across analyses from multiple TAS subsets were combined using the Fisher’s method, and adjusted for multiple hypothesis testing using False Discovery Rate (FDR) procedure (Benjamini and Hochberg 1995).

### Comparison to Drugmatrix signatures

Using gene set enrichment analysis (GSEA) (Subramanian et al. 2005), we compared how well our profiles recapitulated external signatures of carcinogenicity and genotoxicity extracted from the NTP Drugmatrix database (Ganter et al. 2005). The Drugmatrix is a compendium of microarray profiles of short-term chemical exposures in intact rat organs (liver samples used only) and in cell cultures (primary rat hepatocytes). The Drugmatrix-derived signatures were defined as the lists of genes in the Drugmatrix significantly associated with long-term carcinogenicity and genotoxicity. Data processing of the Drugmatrix data is consistent with methods described in Gusenleitner et al. (2014). Gene features were mapped from rat Ensembl gene identifiers to human gene symbols using Biomart (Durinck et al. 2005). Differential expression analysis was conducted using *limma* (Ritchie et al. 2015; Smyth 2005) to identify markers of carcinogenicity and genotoxicity after correcting for the effect of dose and duration of exposure. For each comparison, a list of significant genes was derived using a FDR cutoff of 0.01 and absolute value of log fold change of 0.2, up to a maximum of 300 genes as ranked by FDR. Signatures of carcinogenicity and genotoxicity (direction sensitive: upregulated/downregulated) were derived for three Drugmatrix subsets: liver profiles, cell culture profiles, and low-dose cell culture profiles (< 50μM), the latter consistent with the range of doses used in our experiment. (For detailed gene lists included in the Drugmatrix signatures, see Table S8). These gene signatures were tested for enrichment against our L1000 profiles in various subsets (TAS > 0, 0.2, 0.3, 0.4), using the binary phenotypes of carcinogenicity and genotoxicity and the GSEA method, with empirical p-values estimated based on 10,000 gene-set permutations.

### Comparison with CMap signatures

We performed a systematic comparison of our signatures to those in the CMap database. To this end, we computed the *connectivity score*, a measure of similarity, between pairs of signatures, in this case, between each of our signatures and each of the perturbation signatures in the CMap, which comprises ~1.3 million profiles corresponding to 19,811 drugs and small molecules, and 5,075 molecular (gene-specific knockdown and over-expression) perturbations across 3 to 77 cell lines (Subramanian et al. 2017). The connectivity scores are expressed as percentile values in the [−100, 100] range, wherein a score of 100 represents maximum signature overlap, −100 represents maximum signature reversal and 0 represents lack of concordance between signatures in either direction. Connectivity scores were computed with respect both to individual CMap perturbagens, and to Perturbagen Classes (PCLs), defined as sets of perturbagens with similar MoAs or gene target annotations. Next, we performed differential connectivity analysis with respect to our chemical groups (carcinogens vs. non-carcinogens, genotoxicants vs. non-genotoxicants) using a one-tailed Wilcoxon rank-sum test to test for presence of increased connectivity in the positive class (carcinogenic or genotoxic). These tests were repeated for each TAS-based subset of our data, and false discovery rate (FDR) values were calculated. A minimum mean connectivity score of 60 for the positive class was used to filter out differential connectivity hits with low base connectivity scores.

### Investigation of AhR activation in L1000 profiles

To examine the behavior of AhR-related chemicals included in the study, we tested whether these chemicals exhibit enriched activity of AhR-related gene-sets compiled from independent sources. Lists of chemicals with known AhR activity were identified using multiple AhR-related Tox 21 reporter assays extracted from the tool Tox21 Enricher, or using custom chemical annotation with expert knowledge (referenced as “Sherr_AHR_agonist” in Figure 7A). Lists of AhR target genes were compiled from literature, as annotated in Table S15.

A one-directional weighted Kolmogorov-Smirnov (KS) test was performed to test for the enrichment of “AhR-positive” samples (profiles corresponding to AhR-related chemicals) among the top-ranked profiles sorted by descending AhR geneset activity scores. The activity scores represent the median scores across four individual AhR geneset scores calculated using GSVA.

Profiles corresponding to AhR-related chemicals in the list “Sherr_AHR_agonist” were clustered using the similarity matrix derived from the connectivity scores of the selected profiles (see previous section for the calculation of connectivity scores).

## Results

### TAS analysis and chemical “bioactivity”

We used the *transcriptional activity score* (TAS) as a proxy for chemical bioactivity. Subsequent analyses are based on subsets of profiles at different TAS thresholds (TAS > 0, 0.2, 0.3, 0.4). TAS > 0.2 is the standard cutoff for sufficient bioactivity adopted by the CMap-L1000 workflow (Subramanian et al. 2017), while TAS > 0.3 and TAS > 0.4 represent more stringent thresholds we use to assess the effect of increasing bioactivity on downstream analysis such as classification and gene-set enrichment. While the majority of our profiles have low transcriptional bioactivity, a substantial percent of profiles achieved sufficient TAS. Among 330 chemicals represented across 1972 replicate collapsed profiles, 133 chemicals (40.3%) achieved TAS > 0.2 in at least one dose, 89 chemicals (26.97%) achieved TAS > 0.3 and 63 chemicals (19.09%) achieved TAS > 0.4.

### Chemical dose has a significant effect on transcriptional bioactivity

We performed statistical tests to compare TAS of adjacent dose groups and evaluate how bioactivity is affected by dose. Statistically significantly higher TAS were found when comparing dose rank 3 with rank 2 (p-value < 0.01), rank 4 with 3, rank 5 with 4 and rank 6 with 5 (p-value< 0.001)(Figure 1A). The consistent significance of TAS differences between adjacent dose groups implies that increasing dose is effective at increasing the transcriptional bioactivity of profiles, with the maximum dose used in this experiment yielding the highest range of TAS scores. When binned by TAS range (Figure 1B), the monotonically increasing dose response of TAS is apparent across all bins and stronger for higher TAS ranges.

**Figure 1.**
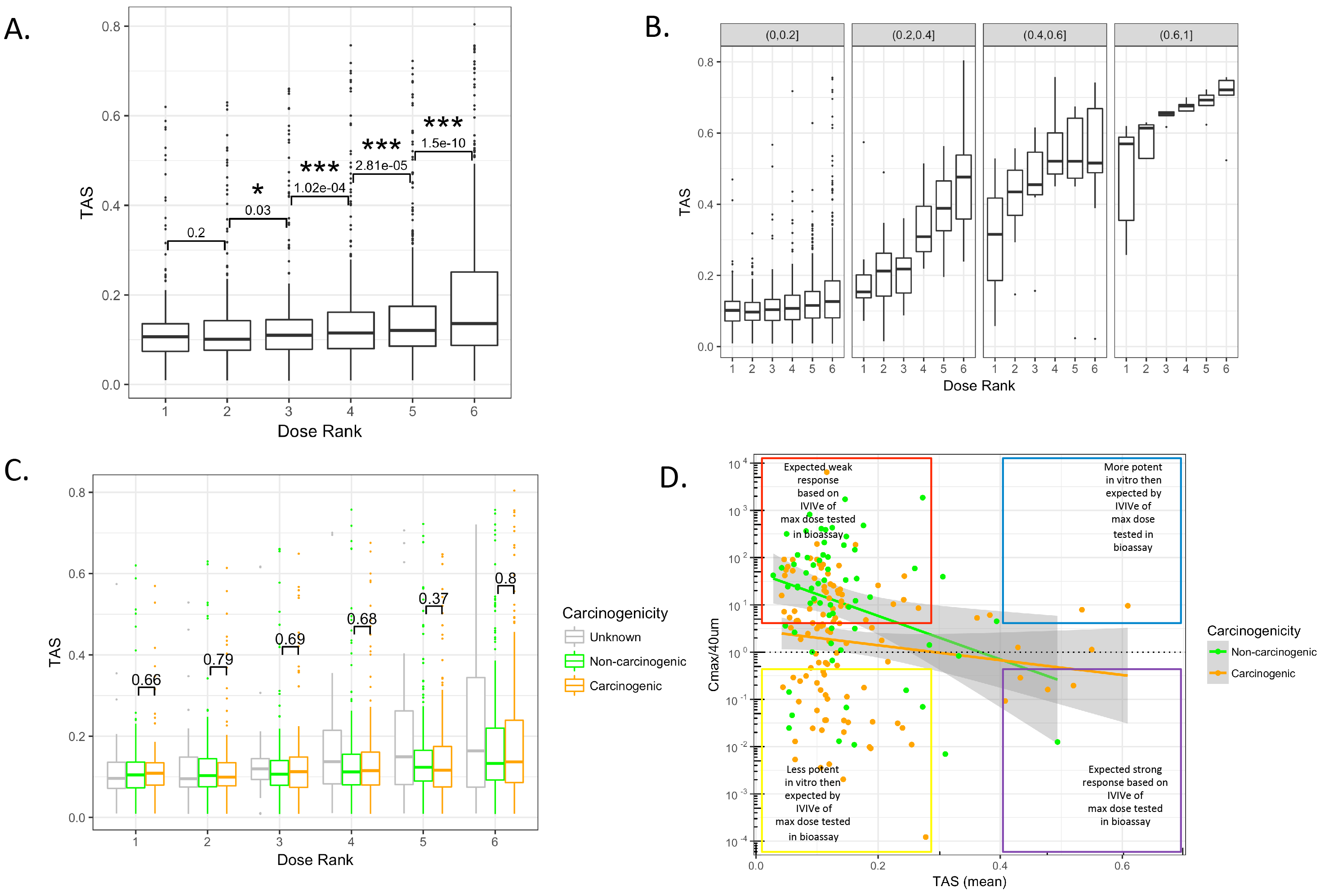
Boxplot of TAS by sample subsets: (A) Boxplot of TAS distributions for each dose level (rank = 1 lowest dose, rank 6 = highest dose). P-values indicate the significance of paired one-sided two-group TAS comparison between adjacent dose groups (* = p< 0.05, ** = p< 0.01, *** = p< 0.001) (see methods). (B) Boxplot of TAS distribution for each dose level, binned by TAS subsets. P-values indicate the significance of paired one-sided two-group TAS comparison between adjacent dose groups (* = p< 0.05, ** = p< 0.01, *** = p< 0.001) within each TAS bin (see methods). (C) Distribution of TAS grouped by chemical carcinogenicity within each dose level. P-values indicate the significance of unpaired one-sided two-group TAS comparison between TAS of carcinogenic chemicals and TAS of non-carcinogenic chemicals within each dose group (* = p< 0.05, ** = p< 0.01, *** = p< 0.001) (see methods). (D) Scatter plot of TAS (mean TAS per chemical) and the ratio of Cmax over maximum in-vitro dose (40uM) (see methods for Cmax calculation).

### Transcriptional bioactivity levels are not associated with carcinogenicity

Next, we evaluated whether the level of a chemical bioactivity as captured by TAS had any association with that chemical’s long-term carcinogenicity or genotoxicity. Remarkably, carcinogenicity showed no effect on TAS in all dose groups (Figure 1C).

On the other hand, genotoxicity showed a marginally significant effect on TAS among profiles with dose rank 1 (lowest dose group) and dose rank 6 (highest dose group) where genotoxic chemicals had significantly higher TAS compared to non-genotoxic chemicals (p-value cutoff > 0.05)(Figure S2).

### In-vitro bioactivity is negatively associated with the rat bioassay dose

The lack of association between TAS and carcinogenicity motivated us to further investigate the relationship between the L1000 doses and the in-vivo doses used in the rodent bioassay. To this end, we tested the association between in-vitro bioactivity (TAS) and the estimated *equivalent in-vitro dose*, Cmax (see Methods), where Cmax represents the estimated in-vitro dose corresponding to the in-vivo dose tested in the rat bioassay. Cmax estimates could be calculated for 183 of the 330 chemicals included in our screen.

Figure 1D shows the mean TAS of profiles for each chemical against the same chemical’s Cmax/40μM (the ratio of estimated equivalent dose to the max in-vitro dose). First, we observe that a substantial number of chemicals have Cmax/40μM values greater than 1, indicating that among these chemicals, higher doses were tested in the rodent bioassay than in our in-vitro assay.

Secondly, we tested the significance of the effect of TAS, carcionogenicity (CARC), and interaction of TAS and CARC on Cmax using regression analysis (see Methods). We found significant effects of TAS (β_TAS_= −4.49, p-value = 0.01), and CARC (β_CARC_ = −1.22, p-value = 0.001) and non-significant effect of the interaction of TAS and CARC (β_T:C_ = 3.1, p-value = 0.16).

As expected, we observed that TAS negatively associates with Cmax for both carcinogens and non-carcinogens. In other words, chemicals that require a low equivalent dose to elicit a carcinogenic response in the rodent bioassay tend to be more transcriptionally active in the in-vitro assay. Interestingly, carcinogenicity also has an effect on Cmax prediction, with non-carcinogens having higher Cmax in general.

### Chemical profiles with aberrant TAS levels can be explained by dose or pharmacological factors

We annotated L1000 chemicals that exhibit unexpected TAS levels, namely, the carcinogenic chemicals with low TAS and non-carcinogenic chemicals with high TAS, to provide potential explanations for the observed TAS behaviors.

We annotated the carcinogenic chemicals with low TAS based on their structural group membership, in-vivo dose requirement for carcinogenicity labeling, and requirements for metabolic activation in HEPG2. Carcinogenic chemicals with low TAS tend to fall in one or more of the following categories: (1) small nitrosamines and other alkylating agents that form DNA adducts but are not adequately recognized by the DNA repair machinery (enriched in yellow box in Figure 1D), (2) require bioactivation by CYP2E1 and other p450s that are not present at high levels in HEPG2 cell culture (also enriched in yellow box in Figure 1D), or (3) require high equivalent In-vitro dose to be carcinogenic, thus likely under-dosed in our in-vitro assay (enriched in red box in Figure 1D).

Among non-carcinogenic chemicals with high TAS, we generally noted low dose used in the rodent bioassays due to toxicity or early deaths at higher doses, e.g., Cyclosporin A (immune suppression), Pyrimethamine and Rhodamine 6G (bone marrow suppression), hexachlorocyclopentadiene (neurotoxicity), Rotenone (mitochondrial effects). Thus, if higher doses were tolerated in rodent bioassays, it is possible that some of these chemicals may elicit a carcinogenic response.

### L1000 profiles with sufficient transcriptional bioactivity accurately predict carcinogenicity and genotoxicity

While a chemical bioactivity level is not predictive of long-term carcinogenicity, the most relevant question is whether a chemical’s bioactivity affects the ability of its expression profile to be predictive of carcinogenicity (and genotoxicity). To answer this question, we built multiple classifiers based on profiles with TAS values within various ranges, and used a random resampling scheme to assess their prediction performance. Datasets corresponding to different TAS ranges were randomly split into train (70%) and test (30%) sets multiple times (n=25), classifiers were built on the train sets, and predictions made on the test sets. The average Area Under the Curve (AUC), sensitivity, and specificity were then estimated over the 25 random resamples. As shown in Figure 2, the prediction AUC improves with higher stringencies of TAS. We achieved the highest predictive accuracy within the most stringent TAS subset (TAS > 0.4), with 72.2±2.7% (mean±se) AUC for prediction of carcinogenicity (Figure 2A), and 82.3±1.6% AUC for prediction of genotoxicity (Figure 2B). These results suggest that short in-vitro gene expression profiles of chemical perturbations, given sufficient transcriptional bioactivity, can accurately predict long-term chemical carcinogenicity and to a greater extent, genotoxicity.

**Figure 2.**
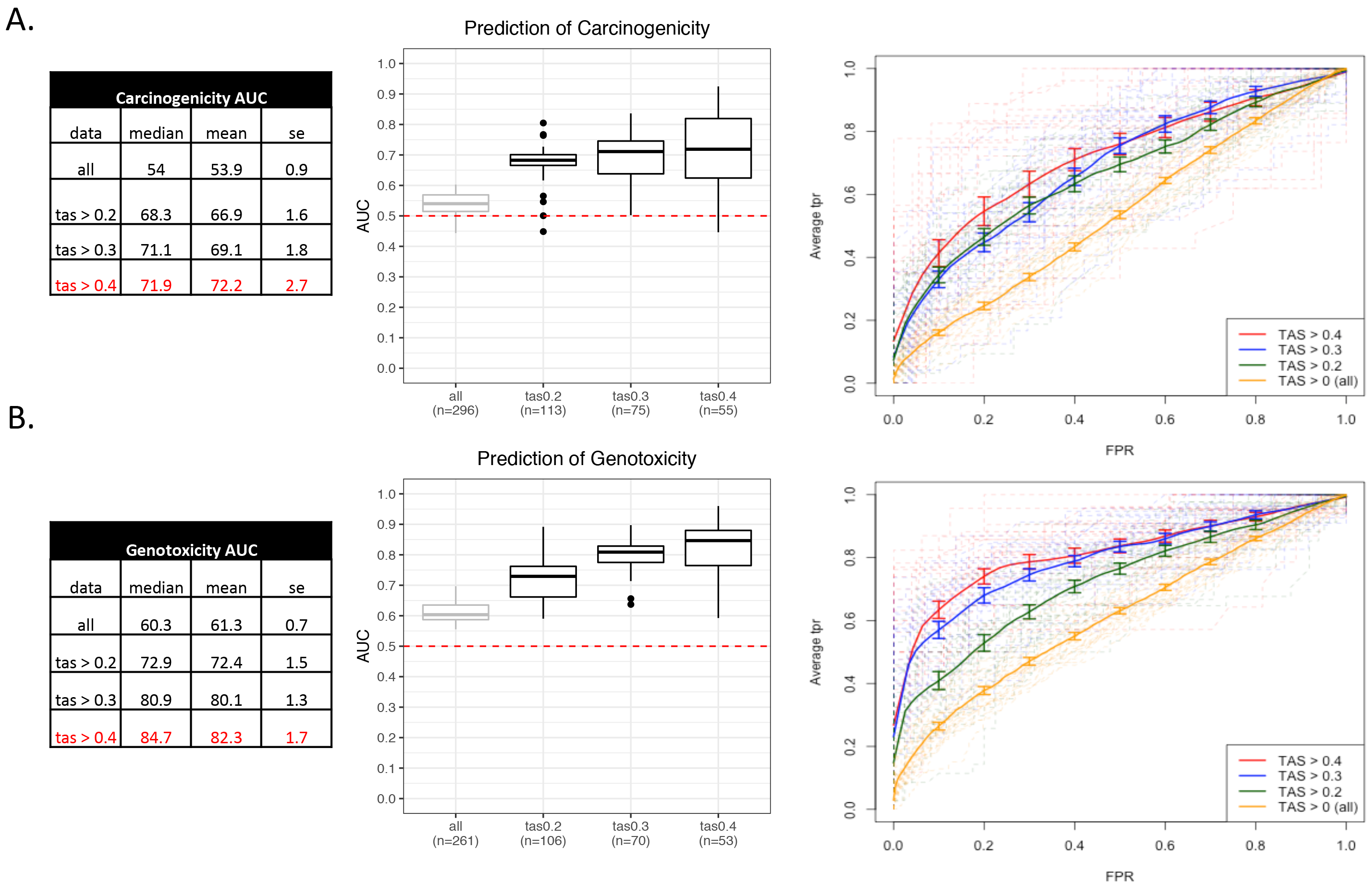
Performance of classifiers in predictive models of (A) carcinogenicity, and (B) genotoxicity, showing, from left to right: summary statistics tables, boxplots of AUC across resamples by TAS subsets, ROC curves of each resample (n = 25) for each TAS subset, and ROC curves aggregating predictions across resamples with points corresponding to thresholds of 0.4, 0.5, and 0.6 for calling binary predicted labels.

### Gene markers for prediction of carcinogenicity and genotoxicity

Final predictive models of carcinogenicity, genotoxicity, and genotoxicity within carcinogens were built using the entire set of profiles with TAS > 0.4. Landmark genes were ranked by variable importance as measured by the mean decrease in Gini coefficient (Table S2) and the top 20 genes for each model were reported in Figure 3. In the carcinogenicity prediction model, top genes include BLCAP, an apoptosis inducing gene, and SESN1, a target of p53 in response to DNA damage and oxidative stress (Figure 3A). Among the top 20 landmark genes for prediction of genotoxicity are pro-apoptotic regulators such as BLCAP and BAX (Figure 3B). Of note, BAX is regulated by p53 and has been shown to be involved in p53-mediated apoptosis, a hallmark of DNA damage response to genotoxic chemical exposure.

**Figure 3.**
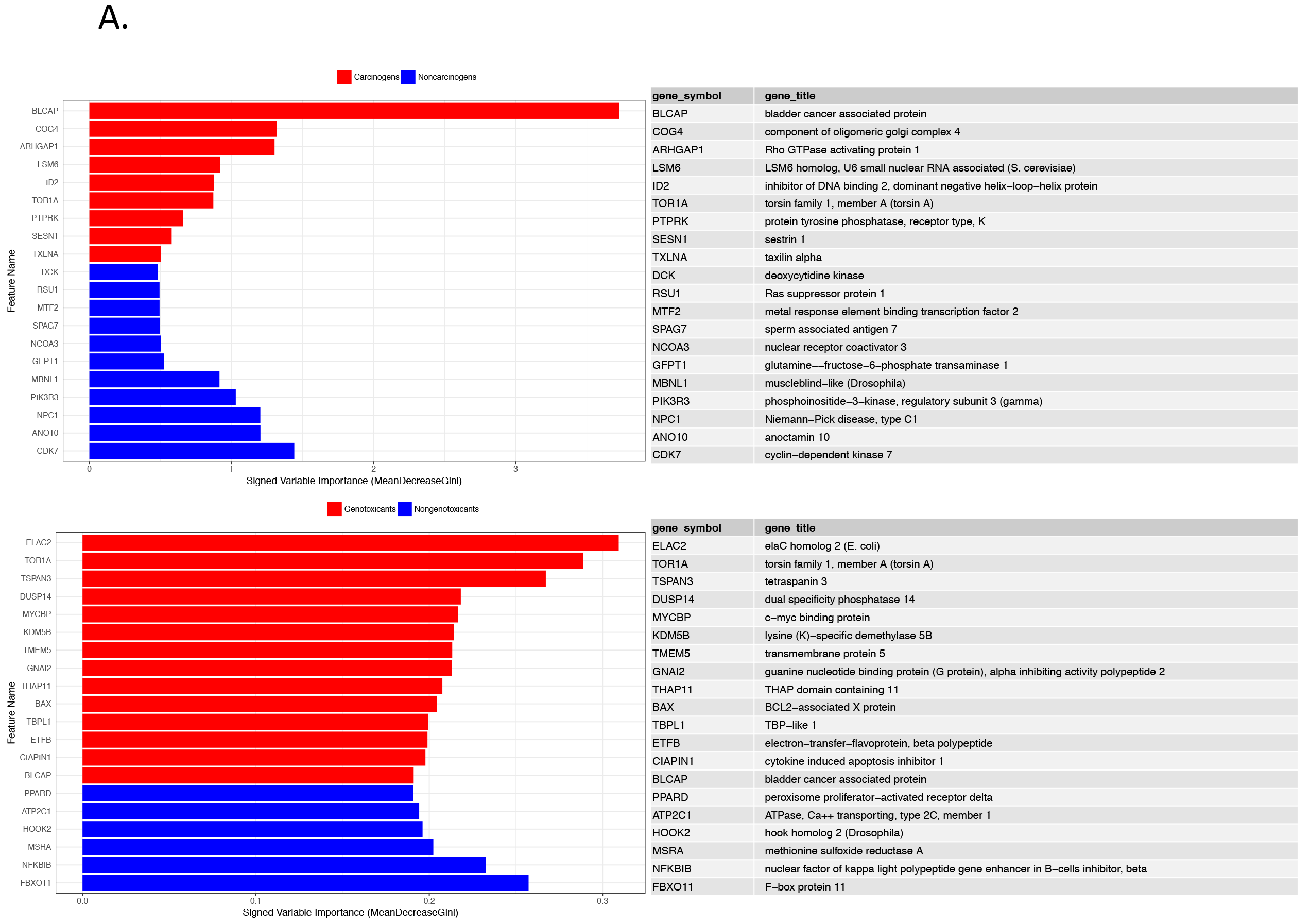
Top features for prediction of carcinogenicity ranked by variable importance in predictive models of (A) carcinogenicity (B) genotoxicity in TAS 0.4 model.

### Final predictions of carcinogenicity and genotoxicity in bioactive profiles

Final predictions of carcinogenicity and genotoxicity were made using a leave-one-chemical-out cross-validation scheme, in which predictive models were trained based on all but one chemical and predictions were made on the profiles of the left-out chemical (see methods). This procedure was repeated for all unique chemicals in profiles with TAS > 0.4 to derive probability measurements of the profile being “Positive” for either carcinogenicity or genotoxicity (see methods) using a probability threshold of 0.5. Prediction probabilities for carcinogenicity and genotoxicity were reported along with the true class labels as the dot colors (Figure 4). From this representation, we observe that predictions tend to be consistent across profiles of varying doses of the same chemical. Several exceptions exist in chemicals whose prediction probabilities were close to 0.5. For example, profiles of Hexachlorocyclopentadiene exposure yield two true negative predictions but one false positive prediction at the highest dose. In addition, prediction probabilities monotonically increasing as a function of dose are observed for some compounds. For example, 3’-Methyl-4-dimethylaminoazobenzene shows increased probability of genotoxicity prediction with increasing dose. However, this pattern is not generalizable to all chemicals. Detailed predictions on TAS>0.4 chemicals are summarized in Table S3.

**Figure 4.**
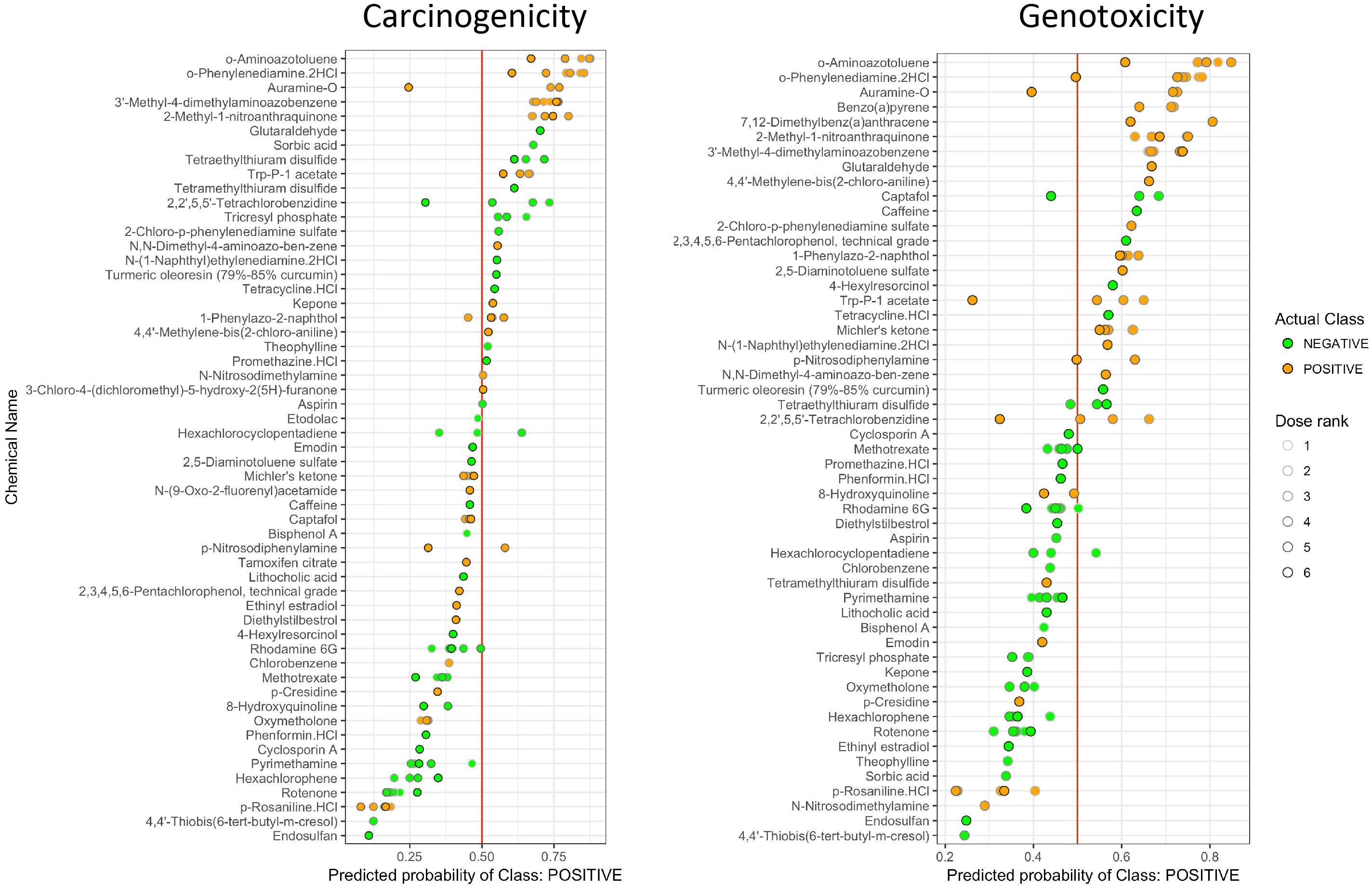
Dot plot of probabilities of predicted classes for hold-out chemicals in the TAS 0.4 subset. Dot fill colors represent actual class labels. Dot outline colors represent dose ranks. X-axis positions of dots represent predicted probability of class “Positive” (carcinogenic in column one or genotoxic in column two).

### Predictions of unlabeled chemicals

Using the final predictive models trained on all profiles with TAS > 0.4, predictions of carcinogenicity and genotoxicity were made for the chemicals without known CPDB annotation (Figure S3, with detailed summary in Table S4 and Table S5). The majority of unlabeled profiles were predicted “Positive” for both carcinogenicity and genotoxicity using a probability threshold of 0.5. This is likely due to bias in chemical selection. Sources of unknown chemicals include chemicals of interest to the Superfund Research Program (likely environmental toxicants), chemicals that were tested for either carcinogenicity or genotoxicity in the CPDB but whose labels cannot be determined, and controversial chemicals in commercial use (triclosan, Glycel). Many profiles have predicted probabilities between 0.5-0.65, indicating low confidence in prediction, potentially attributable to low bioactivity of profiles. When restricting predictions to unlabeled profiles with TAS > 0.4 to be consistent with the subset used for model training, the separation of ranges of prediction probabilities becomes clearer (Figure S3 B, D). The top two ranked predicted carcinogens, benzo(a)pyrene and 7,12-Dimethylbenz(a)anthracene, are two polyaromatic hydrocarbons (PAHs) that have been shown to manifest carcinogenic and genotoxic properties.

The top ranked predicted genotoxicant, indoxyl sulfate, is an endogenous tryptophan metabolite, which has been shown to activate p53 expression through reactive oxygen species (ROS) production and is a source of endogenous oxidative DNA damage (Shimizu et al. 2013). While indoxyl sulfate may not necessarily be considered a genotoxicant as it is a uremic solvent found in low concentrations (1-5μM) in the human serum normally, it activates the AhR, inducing cytochrome P450 enzymes which metabolize other substrates, including mutagenic intermediates. Thus, prediction of indoxyl sulfate as a genotoxicant may be due to transcriptional activation of shared pathways involved in metabolism of genotoxic chemicals. Upon closer inspection of the clustering of AhR ligands in the space of L1000 profiles (Figure 7B), the profiles of indoxyl sulfate perturbations cluster within the group of strong AhR agonists that are mostly known carcinogens or genotoxicants, e.g. benzo(a)pyrene, 7,12-Dimethylbenz(a)anthracene, and TCDD, and distal to the cluster of non-carcinogenic and mainly endogenous AhR ligands.

### Pathway enrichment analysis reveals relevant MoAs of carcinogenicity and genotoxicity

To identify pathway level differences between carcinogens and non-carcinogens, and similarly, between genotoxicants and non-genotoxicants, we performed differential pathway enrichment analysis and ranked pathways according to the significance of their differential enrichment between chemical groups. In accordance with the breakdown of TAS subsets used in classification analysis, and based on the observation that increasing thresholds of TAS yield better classification performance, the differential pathway enrichment analysis was repeated for each of the TAS subsets previously considered (Table S6 and S7). With no TAS threshold (e.g. inclusion of all profiles), only a few pathways are differentially scored between carcinogens and non-carcinogens and between genotoxicants and non-genotoxicants. With increasing thresholds of TAS, the number of significantly expressed pathways increases. At TAS 0.2 and above, the identity of significant pathways becomes more stable, particularly for genotoxicity-related pathways, with many significant pathways shared across TAS > 0.2, 0.3, and 0.4. We derived an aggregated ranking score of differential pathway enrichment by combining p-values across all the TAS subsets (see methods) and included lists of differentially enriched pathways (combined FDR < 0.05) with respect to carcinogenicity in Table S6 and genotoxicity in Table S7.

When comparing carcinogens to non-carcinogens, we observed up-regulation of immune-related pathways (interferon-α/β), cell death (apoptosis induced DNA fragmentation), DNA repair (nucleotide excision repair), transcriptional regulation (RNA polymerase I, II, and III related activity), and cell cycle checkpoints (p53-dependent G1 DNA damage checkpoint), and down-regulation of various metabolism related pathways (phase II conjugation, phase I functionalization, peptide hormone biosynthesis), cell-cell organization and communication (cellcell junction organization, integrin cell surface interactions, tight junction interactions), and G-protein signaling. Among genotoxicants compared to non-genotoxicants, upregulated pathways include DNA repair (nucleotide excision repair, formation of incision complex in GG-NER), AKT signaling, programmed cell death, G1/S DNA damage checkpoints, innate immune response (interferon signaling, toll-like receptor signaling). Down-regulated pathways include xenobiotic metabolism (phase I and phase II metabolism), peptide hormone biosynthesis, cell-cell organization and cell-cell communication, innate immune response (complement cascade), and various hemostasis and metabolism related pathways.

From the differentially scored pathways of carcinogenicity (Table S6) and genotoxicity (Table S7), we identified a reduced set consisting of the top 40 up-regulated and top 40 down-regulated pathways with Reactome categories as ordered by the aggregated rankings, and visualized their enrichment scores across profiles with TAS > 0.2 in Figure 5A (top pathways differentially enriched with respect to carcinogenicity) and Figure 5B (genotoxicity). Hierarchical clustering of samples reveals loose stratification by carcinogenicity status (carcinogens in orange) and stronger stratification by genotoxicity status (genotoxicants in purple).

**Figure 5.**
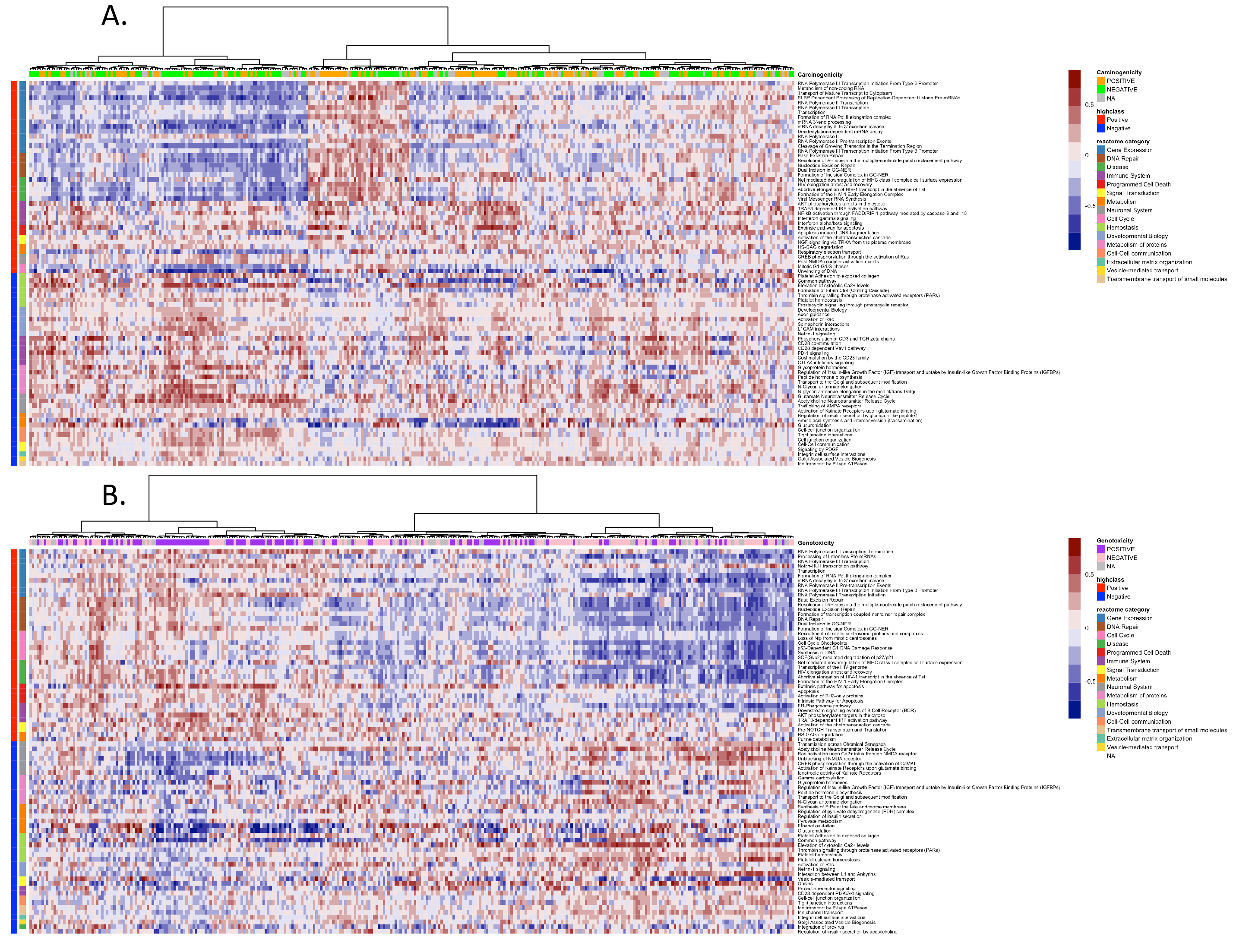
Heatmap of pathway enrichment scores (GSVA) for top 40 upregulated and downregulated differential pathways of carcinogenicity (A) and genotoxicity (B) for profiles with TAS > 0.2. Columns are clustered using the ward method with euclidean distances. Rows are ordered by the frequency of the pathway categories among the top 40 (direction sensitive).

### L1000 signatures of carcinogenicity and genotoxicity are consistent with signatures from low-dose cell culture in Drugmatrix

We tested for enrichment of the Drugmatrix-derived signatures of carcinogenicity and genotoxicity against our L1000-based differential signatures of carcinogenicity and genotoxicity (Table S9). Significant similarities were observed between signatures derived from Drugmatrix low dose rat primary hepatocyte cell cultures and our L1000 profiles. For example, the signature of up-regulated genes in response to low-dose carcinogens in cell cultures (UP_CARC_CELL_LOWDOSE) was enriched in the L1000-profiled carcinogen subsets at TAS > 0.4, 0.3, and 0.2 (FDR<0.05). Conversely, the signature of down-regulated genes in response to low-dose carcinogens in cell cultures (DN_CARC_CELL_LOWDOSE) was enriched in the L1000-profiled non-carcinogen subsets at TAS > 0.2 and 0 (FDR<0.05). Similarly, signature of genotoxicants in the Drugmatrix cell cultures (“UP_GTX_CELL_LOWDOSE”) was enriched in the L1000-profiled genotoxicant subsets at TAS > 0.4, 0.3, and 0. When repeating the analysis for signatures derived from *all* Drugmatrix cell culture profiles (including high doses), signatures of genotoxicity were mostly directionally consistent with L1000 profiles, but signatures of carcinogenicity were inconsistent, and in fact sometimes behaving in the opposite direction (e.g. Drugmatrix signature “UP_CARC_CELL” has enrichment among non-carcinogens in L1000 TAS > 0.4 profiles). This inconsistency may be explained by the presence of extremely high doses used for some chemicals in generating the Drugmatrix cell culture profiles. For reference, the mean dose in Drugmatrix cell culture profiles is ~3,000uM and the max dose is 180mM. In contrast, the max dose among L1000 profiles is 40uM.

Next, we compared signatures derived from the Drugmatrix in-vivo rat liver profiles to the L1000 profiles. For carcinogenicity, the signature of down-regulated genes in response to carcinogens (“DN_CARC_LIVER”) is correctly enriched among non-carcinogens in L1000 TAS < 0.4, 0.3, 0.2 and 0 with FDR < 0.05. Similarly, the signature of up-regulated genes in response to carcinogens (“UP_CARC_LIVER”) is marginally enriched among L1000 TAS < 0.4 (FDR = 0.06), and TAS < 0.3 (FDR = 0.09) carcinogens. On the other hand, the signatures of genotoxicity are largely not enriched in the right direction (e.g., “DN_GTX_LIVER” shows enrichment among genotoxicants of TAS 0.4).

To rule out the possibility that the observed signatures’ inconsistency is due to platform differences – since the Drugmatrix data is microarray-based while our data is generated from the L1000 – we compared Drugmatrix cell culture to Drugmatrix liver signatures of genotoxicity (both microarray based). We found that the downregulated genotoxicant signature in liver is also behaving in the opposite direction than in cell culture (Table S10). This finding suggests that the signatures’ inconsistency between liver and cell line is likely due to differences between in-vitro and in-vivo responses to exposure rather than to differences in the profiling platform. Upon detailed inspection of the Drugmatrix liver signatures, we identified an enrichment of genes relating to metabolism in both the up and down regulated gene signatures (lipid metabolism, cholesterol biosynthesis, Phase I metabolism in “UP_GTX_LIVER”, amino acid metabolism, fatty acid metabolism in “DN_GTX_LIVER”), supporting the conclusion that there may be substantial differences between metabolic activities in in-vitro and in-vivo exposures (Figure S4).

In summary, L1000 derived signatures of carcinogenicity and genotoxicity are concordant with Drugmatrix low dose cell culture signatures, but inconsistent with Drugmatrix liver signatures, with the differences largely driven by discrepancies in the expression of certain metabolism-related genes between in-vitro and in-vivo exposures.

### L1000 signatures of carcinogenicity and genotoxicity capture biologically relevant MoAs of several drug classes in the CMap

The availability of the CMap offers the opportunity to compare our profiles to a much larger database of pharmacologically annotated signatures and allows us to predict MoAs or pharmacological properties based on signature similarity. To this end, we first computed the similarity of our signatures to each signature in the CMap. We then identified the CMap signatures that show significant difference in connectivity to carcinogens and non-carcinogens, and to genotoxicants and non-genotoxicants. The top CMap hits are summarized at the level of Perturbagen Classes (PCLs) in Table S11 (carcinogenicity) and Table S12 (genotoxicity) and visualized in Figure 6, and at the level of individual chemical perturbations in Table S13 (carcinogenicity) and Table S14 (genotoxicity).

**Figure 6.**
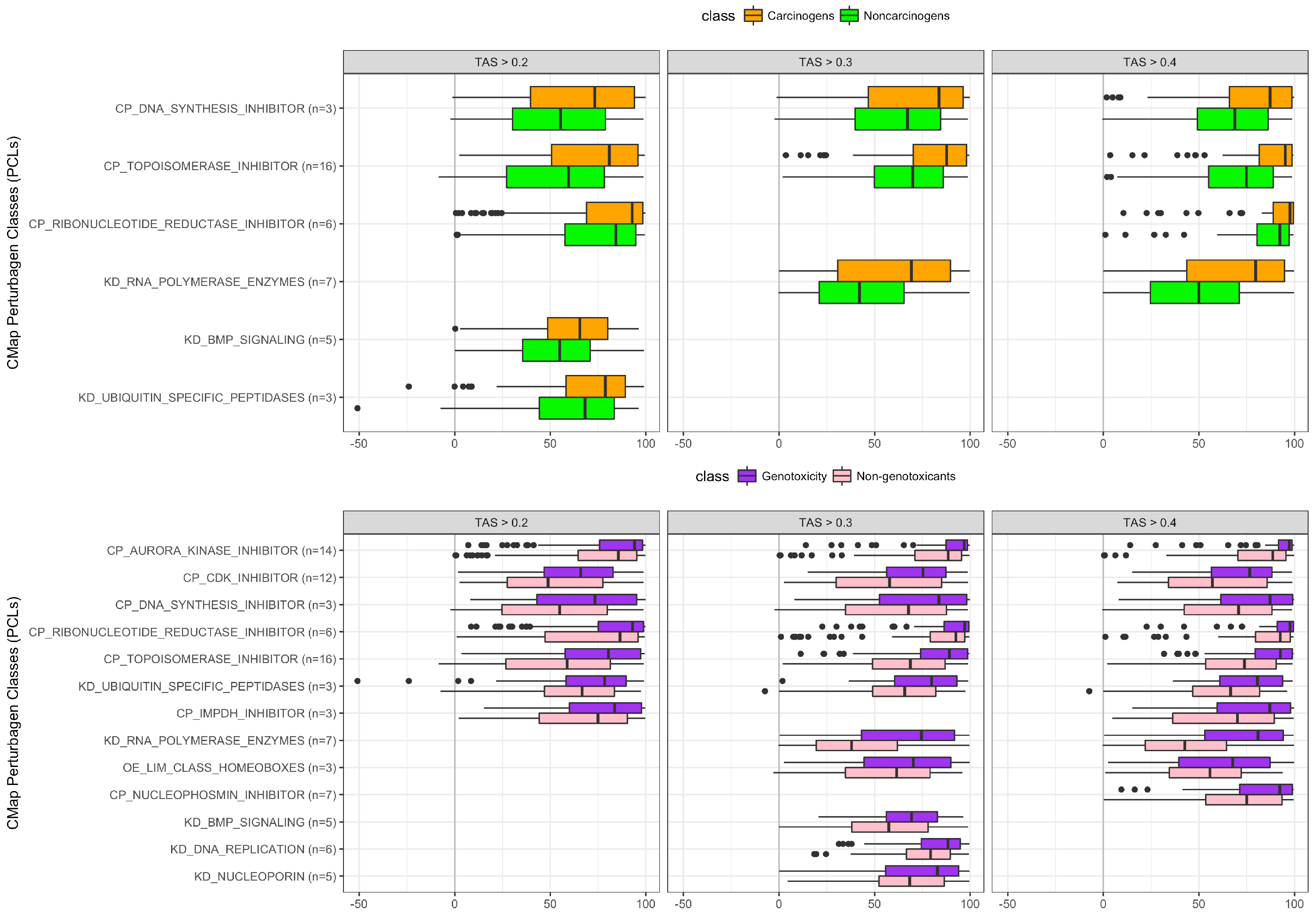
Connectivity scores of top CMap Perturbagen Classes to Carcinogens vs. Non-carcinogens, Genotoxicants vs. Non-genotoxicants.

Focusing on the significantly differential PCLs across all TAS subsets (TAS > 0.2, 0.3, 0.4), we found that carcinogens, compared to non-carcinogens, are significantly more connected to drug classes consisting of topoisomerase inhibitors, DNA synthesis inhibitors, and ribonucleotide reductase. Genotoxicants, compared to non-genotoxicants, are significantly more connected to the three aforementioned drug classes, as well as to CDK inhibitors, aurora kinase inhibitors, and ubiquitin specific peptidases (Figure 6).

Topoisomerase inhibitors, a specific class of DNA synthesis inhibitors, are most recognized as chemotherapeutic drugs that preferentially inhibit the topoisomerase enzymes (commonly topoisomerase I or II) in cancer cells to slow their rate of replication. Topoisomerase I or II introduce single- or double-strand DNA breaks in cells undergoing replication, and form topoisomerase-DNA complexes. Most topoisomerase inhibitors function by trapping these complexes, leading to increased strand breaks but incomplete DNA replication, subsequently provoking DNA damage response and DNA repair (Pommier 2006; Pommier 2013; Wang et al. 2002). Thus, DNA damage response induced by topoisomerase inhibitors is expected to mimic the response to genotoxic carcinogens.

Other relevant PCLs also exhibit shared MoAs with carcinogens and genotoxicants. Aurora kinase inhibitors play a major role in cell cycle regulation through the induction of G1 arrest and apoptosis (Bavetsias and Linardopoulos 2015). Ubiquitin specific peptidases, specifically USP24, have been shown to play a role in DNA damage response (Zhang and Gong 2016).

### L1000 gene expression profiles of AhR agonists capture independently defined AhR-mediated responses and identify sub-clusters based on receptor binding strength and toxicity

Carcinogens and genotoxicants are often recognized by receptors such as the aryl hydrocarbon receptor (AhR). For example, TCDD, a potent toxicant, and more generally members of a class of halogenated planar hydrocarbons are mediated through AhR. In addition to its role in inducing the toxic effects of planar halogenated hydrocarbons and polycyclic aromatic hydrocarbons (PAH), AhR exhibits endogenous functions such as regulating expression of stem cell-associated genes (Stanford et al. 2016; Wang et al. 2010), T cell differentiation (Apetoh et al. 2010; Gandhi et al. 2010), and amino acid tryptophan metabolism (Cheng et al. 2015; Hubbard et al. 2015).

Given that the AhR is an important mediator of many toxicants with strong representation in our dataset, we sought to investigate the behavior of AhR-activated chemicals in terms of gene expression of known AhR gene targets, and the similarity of profiles among sub-groups of AhR agonists.

As shown in Figure 7A, we found that the L1000 profiles exhibit consistent enrichment of AhR-related gene-set activity among chemicals labeled as AhR-active in several Tox21 reporter assays, namely, HTS_ACTIVE.agonism_AhR (p-value: 2.9e-7), HTS_ACTIVE.cytotoxicity_AhR/agonism (pvalue: 1.5e-4) and TOXCAST.ATG_Ahr_CIS_up (p-value: 0.006). This finding validates the ability of unbiased gene expression profiling to accurately capture endpoints from more specific and targeted assays such as those in the Tox21 library.

**Figure 7.**
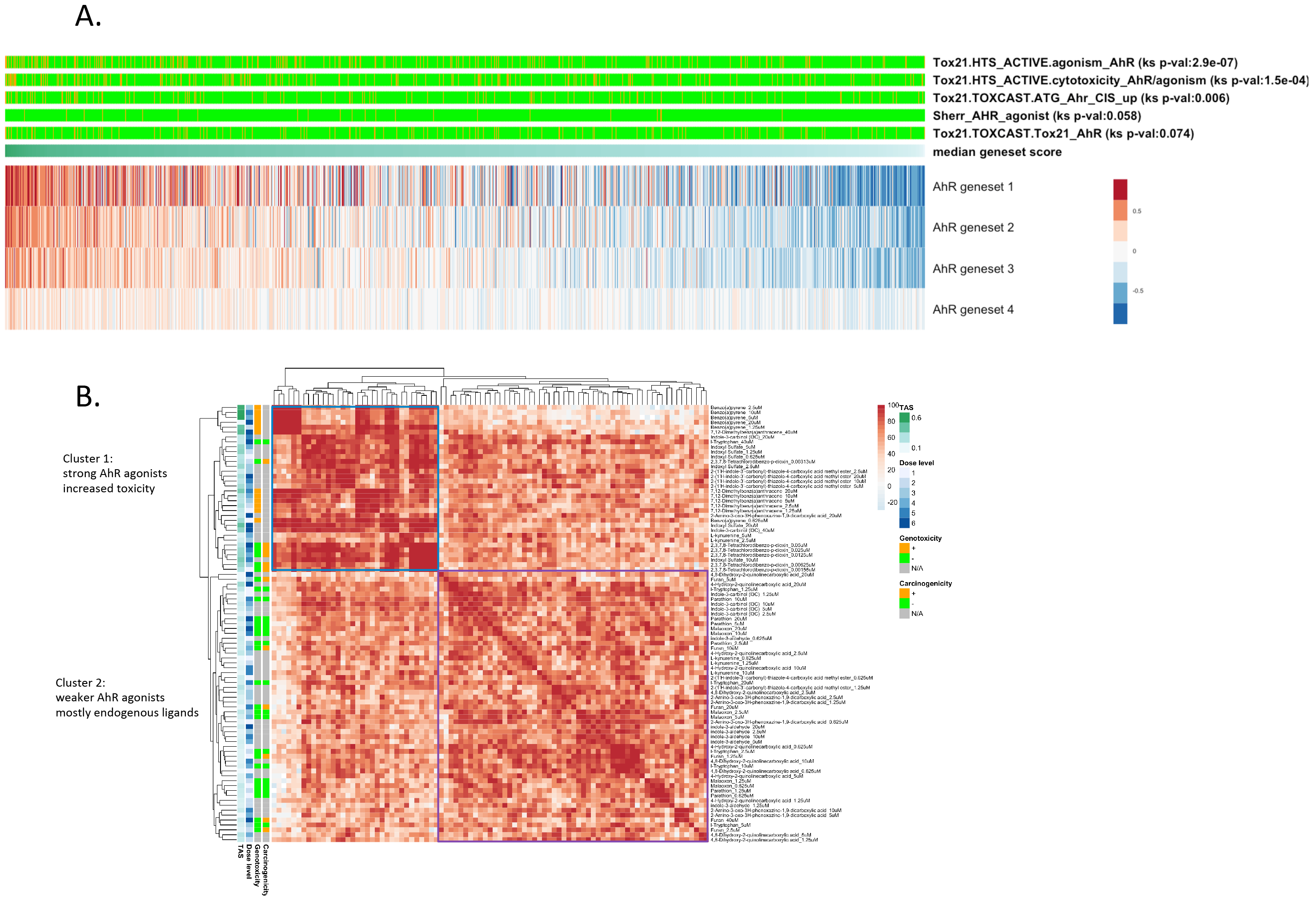
Investigation of profiles of AhR related chemical perturbations: (A) Profiles with AhR activity ranked by median geneset scores of AhR target gene lists. (B) AhR-related profiles clustered by connectivity scores.

Furthermore, we identified sub-clusters of AhR-related chemicals based on the similarity of their gene expression profiles as measured by the connectivity scores and found two functionally distinct classes (Figure 7B). Cluster 1 is enriched for profiles of strong exogenous AhR ligands, most with potent toxic effects (benzo(a) pyrene, 7, 12-Dimethylbenz(a) anthracene, TCDD). It is not surprising that many of these chemicals also have high in-vitro transcriptional bioactivity (high TAS). Cluster 2 contains endogenous AhR ligands (l-kynurenine, indole-3-carbonyl, kynurenic acid, xanthurenic acid, and cinnabarinic acid). Since l-tryptophan is not an AHR ligand, its presence in this latter group suggests that it is metabolized to one of the kynurenine pathway metabolites that are AhR ligands (l-kynurenine, kynurenic acid, xanthurenic acid, and cinnabarinic acid).

These results show promise for our platform to be used not only as a general predictor of general phenotypes such as AhR receptor activation, but also to distinguish, with finer granularity, classes of AhR agonists.

### Carcinogenome Portal – a framework for data query and visualization

All data described in this manuscript are available for public access. Data processed under the standard CMap-L1000 pipeline are available under https://clue.io/data/CRCGN_ABC. To facilitate the interactive querying of the downstream analysis results produced by this study, we developed a web portal (https://carcinogenome.org/HEPG2). The query and visualization functionalities supported by the portal include differential expression, gene-set enrichment, and connectivity analysis against CMap signatures. This interface supports both marker-centered (genes, pathways, CMap signatures) and chemical-centered queries. For instance, one can ask “what gene markers and pathways are regulated by perturbation with Bisphenol A?”, “what are the top chemicals that up-regulate a particular gene or pathway of interest?”, or “which CMap chemicals or chemical groups are most similar to the profiles of perturbation with Bisphenol A in this project?”. In addition, the portal supports bulk query and visualization of groups of perturbations in the form of heatmaps.

## Discussion

### Prediction of carcinogenicity and genotoxicity

The results from the prediction of carcinogenicity and genotoxicity experiments provide strong evidence that transcriptional bioactivity as captured by TAS has a high impact on the classifier performance. That is, while absolute levels of bioactivity are not associated with carcinogenicity, a sufficiently high bioactivity is necessary to elicit enough transcriptional signal to use a chemical’s expression profile for carcinogenicity prediction. Thus, when limiting to profiles with high TAS, the performance of our predictive models drastically improves. Among highly bioactive profiles (TAS>0.4), our classifiers yielded mean AUC of 72.2% for prediction of carcinogenicity (Figure 2A), and 82.3% for prediction of genotoxicity (Figure 2B). To boost the effective sample size used in classification, we outline the following dose selection strategy for improving bioactivity of in-vitro gene expression profiles.

### in-vitro dose recommendation

The selection of doses in short-term acute exposures for prediction of long-term in-vivo phenotypes is a challenging task. In this experiment, we chose to adopt a standard 6-dose titration, starting from 40μM or 20μM depending on source of chemicals. The sole exception to the standard dosing was TCDD, whose starting concentration is 50nM due to its extreme potency. The choice of standard dosing was made for a couple of reasons: 1) lack of commercial availability of certain chemicals at higher stock concentrations; 2) scarcity of in-vitro dose recommendations from publicly available data, e.g., dose recommendations derived from MTT assays; and 3) cost efficiency of standardized dosing using the L1000 platform.

One alternative dosing scheme is to determine unique doses for each chemical using the MTT assay. For instance, a previous study of genotoxicity prediction based on in-vitro experiments selected doses based on a MTT assay resulting in 80% viability at 72h incubation, or maximum dose of 2mM in the case of lack of cytotoxicity (Magkoufopoulou et al. 2012). Some chemicals used in that study were administered at doses that vastly exceeded the 40μM or 20μM dose limit adopted in our experimental setup. Furthermore, the lack of plateau effect in dose response as a function of TAS (proxy for bioactivity) suggests that doses exceeding the 40μM or 20μM threshold may indeed yield profiles with higher bioactivity and increase the power to detect gene and pathway markers for prediction of carcinogenicity and genotoxicity without experiencing saturation effects (response plateauing) or excessive cell death. Although standardizing dosage across chemicals was the logistically and cost-effective solution for this experiment, going forward, MTT assays are highly recommended for maximizing biological signal across transcriptional profiles.

Estimation of the appropriate in-vitro dose from toxicokinetic modeling of the in-vivo doses tested in animal bioassays, when available, is another viable alternative, as shown in Figure 1D and associated discussion.

### Acute vs. chronic response

Through analysis of transcriptional activity scores between carcinogens and non-carcinogens (Figure 1C), we observed that long-term carcinogenicity, as established from long-term in-vivo rodent studies, has no effect on transcriptional bioactivity in our short-term assay (Figure 2A). This observation indicates that bioactivity as defined by TAS at less than 40μM is not associated with carcinogenicity, and consequently, a short-term chemical perturbation with minimal transcriptional response cannot be assumed “safe”.

While TAS alone is not predictive of carcinogenicity, it was instrumental to the selection of those compounds with sufficient bioactivity to allow us to build an accurate gene expression-based classifier of carcinogenicity (up to 72.2% AUC), and to capture certain MoAs of carcinogenicity. Overall, we observe a stronger signal of genotoxicity compared to carcinogenicity, which is to be expected, as the latter is a more heterogeneous phenotype and thus harder to capture as a binary distinction; this is evidenced by the higher accuracy of the genotoxicity classifier (82.3%) as well as the by the higher TAS among genotoxicants compared to non-genotoxicants.

### Challenges and future developments

This experiment aims to accelerate short-term in-vitro testing approaches to predict long-term chemical carcinogenicity. We have shown that short-term in-vitro gene expression profiling is not only capable of accurate prediction of carcinogenicity and genotoxicity, but also useful for characterizing certain mechanisms of carcinogenic response, particularly DNA damage and repair, and changes in cell cycle and cell-cell organization and communication. Other general biological processes that may be relevant for carcinogenic response, including inflammatory response, immune dysfunction, metabolic disruption and endocrine disruption, require further investigation in other in-vitro contexts.

The choice of HEPG2 as our primary cell line model was driven by the abundance of chemical annotations for liver carcinogenicity and the appropriateness of HEPG2 for the study of liver toxicity. However, there are limitations in its use.

Firstly, the expression of genes involved in phase I and phase II metabolism vary between passages and results relating to xenobiotic metabolism may be difficult to determine (Soldatow et al. 2013); this is also seen in the comparison of our genotoxicity-related signatures to Drugmatrix liver signatures. One potential contribution to this effect is the relevantly low bioactivation capacity in HEPG2 compared to in-vivo. It should be noted that, alternatively, the hepatoma cell line, HepaRG, which has a liver-like bioactivation, could be used as an in-vitro liver model for studying carcinogens and genotoxicants. One study has shown that while HEPG2 performs better in discriminating signatures between genotoxic and non-genotoxic carcinogens, HepaRG is a more suitable in-vitro liver model for biological interpretation of effects of chemical exposures (Jennen et al. 2010).

Secondly, since HEPG2 is a cancer cell line, the exposures of carcinogens in this line may show differences as compared to a non-transformed cell line. For the purpose of predictive modeling, these cell line-specific nuances may be overlooked as long as the performance of the classifier is adequate.

While liver carcinogenicity prediction was the adverse phenotype of choice for this study, this experiment provided us with many valuable insights to facilitate future experiments, including logistics of procurement of large chemical panels, chemical and dose selection for tissue specific carcinogenicity. It also sets the stage for in-vitro based exposure studies of additional adverse phenotypes. For instance, we initiated the in-vitro screening of mammary gland carcinogenicity through the use of a non-tumorigenic human mammary epithelial cell line, MCF10A and p53-deficient MCF10A. The experimental and computational pipeline we established, paired with the cost-effective technology we used for chemical exposure and gene expression profiling, paves the way for the screening of large chemical panels for exposure-based experiments in other organ and disease or adverse outcome specific contexts.

## Conclusions

Long term tests for chemical carcinogens based on epidemiology and rat studies are expensive and time consuming and are not feasible for scaling to a large number of chemicals. In this study, we detailed a high-throughput gene expression profiling of more than 300 liver carcinogens and non-carcinogens in a short term in-vitro exposure model. These gene expression profiles, given sufficient transcriptional bioactivity, are capable of accurate prediction of longterm carcinogenicity and even more accurate prediction of genotoxicity. Pathway enrichment analysis revealed similarities between pathway level response captured by the short term in-vitro exposures and known MoAs of carcinogenesis, particularly genotoxic mechanisms such as DNA damage and repair.

## References

American Cancer Society. (2017). Cancer Facts & Figures 2017. Cancer Facts & Figures 2017, 1. doi:10.1097/01.NNR.0000289503.22414.79

Anand, P., Kunnumakara, A. B., Sundaram, C., Harikumar, K. B., Tharakan, S. T., Lai, O. S., … Aggarwal, B. B. (2008). Cancer is a preventable disease that requires major lifestyle changes. Pharmaceutical Research. doi:10.1007/s11095-008-9661-9

Apetoh, L., Quintana, F. J., Pot, C., Joller, N., Xiao, S., Kumar, D., … Kuchroo, V. K. (2010). The aryl hydrocarbon receptor interacts with c-Maf to promote the differentiation of type 1 regulatory T cells induced by IL-27. Nature Immunology, 11(9), 854–861. doi:10.1038/ni.1912

Bavetsias, V., & Linardopoulos, S. (2015). Aurora Kinase Inhibitors: Current Status and Outlook. Frontiers in Oncology, 5. doi:10.3389/fonc.2015.00278

Benjamini, Y., & Hochberg, Y. (1995). Controlling the false discovery rate: a practical and powerful approach to multiple testing. Journal of the Royal Statistical Society B. doi:10.2307/2346101

Bucher, J. R., & Portier, C. (2004). Human carcinogenic risk evaluation, part V: The National Toxicology Program vision for assesing the human carcinogenic hazard of chemicals. Toxicological Sciences, 82(2), 363–366. doi:10.1093/toxsci/kfh293

Cheng, Y., Jin, U.-H., Allred, C. D., Jayaraman, A., Chapkin, R. S., & Safe, S. (2015). Aryl Hydrocarbon Receptor Activity of Tryptophan Metabolites in Young Adult Mouse Colonocytes. Drug Metabolism and Disposition: The Biological Fate of Chemicals, 43(10), 1536–43. doi:10.1124/dmd.115.063677

Croft, D., Mundo, A., Haw, R., & Milacic, M. (2014). The Reactome pathway knowledgebase. Nucleic Acids, 42(D1), D472–D477. doi:10.1093/nar/gkt1102

Durinck, S., Moreau, Y., Kasprzyk, A., Davis, S., De Moor, B., Brazma, A., & Huber, W. (2005). BioMart and Bioconductor: A powerful link between biological databases and microarray data analysis. Bioinformatics, 21(16), 3439–3440. doi:10.1093/bioinformatics/bti525

Eichner, J., Kossler, N., Wrzodek, C., Kalkuhl, A., Bach Toft, D., Ostenfeldt, N., … Zell, A. (2013). A Toxicogenomic Approach for the Prediction of Murine Hepatocarcinogenesis Using Ensemble Feature Selection. PLoS ONE, 8(9). doi:10.1371/journal.pone.0073938

Ellinger-Ziegelbauer, H., Gmuender, H., Bandenburg, A., & Ahr, H. J. (2008). Prediction of a carcinogenic potential of rat hepatocarcinogens using toxicogenomics analysis of short-term in vivo studies. Mutation Research - Fundamental and Molecular Mechanisms of Mutagenesis, 637(1-2), 23–39. doi:10.1016/j.mrfmmm.2007.06.010

Fabregat, A., Sidiropoulos, K., Garapati, P., Gillespie, M., Hausmann, K., Haw, R., … D’Eustachio, P. (2016). The reactome pathway knowledgebase. Nucleic Acids Research, 44(D1), D481–D487. doi:10.1093/nar/gkv1351

Fitzpatrick, R. B. (2008). CPDB: Carcinogenic potency database. Medical Reference Services Quarterly, 27(3), 303–311. doi:10.1080/02763860802198895

Gandhi, R., Kumar, D., Burns, E. J., Nadeau, M., Dake, B., Laroni, A., … Quintana, F. J. (2010). Activation of the aryl hydrocarbon receptor induces human type 1 regulatory T cell-like and Foxp3+ regulatory T cells. Nature Immunology, 11(9), 846–853. doi:10.1038/ni.1915

Ganter, B., Snyder, R. D., Halbert, D. N., & Lee, M. D. (2006). Toxicogenomics in drug discovery and development: mechanistic analysis of compound/class-dependent effects using the DrugMatrix database. Pharmacogenomics, 7(7), 1025–44. doi:10.2217/14622416.7.7.1025

Gold, L. S., Manley, N. B., Slone, T. H., Rohrbach, L., & Garfinkel, G. B. (2005). Supplement to the Carcinogenic Potency Database (CPDB): Results of animal bioassays published in the general literature through 1997 and by the National Toxicology Program in 1997-1998. Toxicological Sciences. doi:10.1093/toxsci/kfi161

Gusenleitner, D., Auerbach, S. S., Melia, T., Gómez, H. F., Sherr, D. H., & Monti, S. (2014). Genomic models of short-term exposure accurately predict long-term chemical carcinogenicity and identify putative mechanisms of action. PLoS ONE, 9(7). doi:10.1371/journal.pone.0102579

Hänzelmann, S., Castelo, R., & Guinney, J. (2013). GSVA: gene set variation analysis for microarray and RNA-Seq data. BMC Bioinformatics, 14(1), 7. doi:10.1186/1471-2105-14-7

Hubbard, T. D., Murray, I. a, & Perdew, G. H. (2015a). Indole and Tryptophan Metabolism: Endogenous and Dietary Routes to Ah Receptor Activation. Drug Metabolism and Disposition, 43(10), 1522–1535. doi:10.1124/dmd.115.064246

Hubbard, T. D., Murray, I. a, & Perdew, G. H. (2015b). Indole and Tryptophan Metabolism: Endogenous and Dietary Routes to Ah Receptor Activation. Drug Metabolism and Disposition, 43(10), 1522–1535. doi:10.1124/dmd.115.064246

Huff, J., Jacobson, M. F., & Davis, D. L. (2008). The limits of two-year bioassay exposure regimens for identifying chemical carcinogens. Environmental Health Perspectives. doi:10.1289/ehp.10716

Jennen, D. G. J., Magkoufopoulou, C., Ketelslegers, H. B., van Herwijnen, M. H. M., Kleinjans, J. C. S., & van Delft, J. H. M. (2010). Comparison of HepG2 and HepaRG by whole-genome gene expression analysis for the purpose of chemical hazard identification. Toxicological Sciences, 115(1), 66–79. doi:10.1093/toxsci/kfq026

Judson, R. S., Houck, K. A., Kavlock, R. J., Knudsen, T. B., Martin, M. T., Mortensen, H. M., … Dix, D. J. (2010). In vitro screening of environmental chemicals for targeted testing prioritization: The toxcast project. Environmental Health Perspectives, 118(4), 485–492. doi:10.1289/ehp.0901392

Kleinstreuer, N. C., Dix, D. J., Houck, K. A., Kavlock, R. J., Knudsen, T. B., Martin, M. T., … Judson, R. S. (2013). In vitro perturbations of targets in cancer hallmark processes predict rodent chemical carcinogenesis. Toxicological Sciences, 131(1), 40–55. doi:10.1093/toxsci/kfs285

Kossler, N., Matheis, K. A., Ostenfeldt, N., Toft, D. B., Dhalluin, S., Deschl, U., & Kalkuhl, A. (2015). Identification of specific mRNA signatures as fingerprints for carcinogenesis in mice induced by genotoxic and nongenotoxic hepatocarcinogens. Toxicological Sciences, 143(2), 277–295. doi:10.1093/toxsci/kfu248

Kuhn, M., Wing, J., Weston, S., Williams, A., Keefer, C., & Engelhardt, A. (2012). Caret: Classification and Regression Training. https://Cran.R-Project.Org/Package=Caret. doi:10.1053/j.sodo.2009.03.002

Liberzon, A., Subramanian, A., Pinchback, R., Thorvaldsdóttir, H., Tamayo, P., & Mesirov, J. P. (2011). Molecular signatures database (MSigDB) 3.0. Bioinformatics, 27(12), 1739–1740. doi:10.1093/bioinformatics/btr260

Magkoufopoulou, C., Claessen, S. M. H., Tsamou, M., Jennen, D. G. J., Kleinjans, J. C. S., & Van delft, J. H. M. (2012). A transcriptomics-based in vitro assay for predicting chemical genotoxicity in vivo. Carcinogenesis, 33(7), 1421–1429. doi:10.1093/carcin/bgs182

Noguchi, S., Saito, A., Horie, M., Mikami, Y., Suzuki, H. I., Morishita, Y., … Nagase, T. (2014). An Integrative Analysis of the Tumorigenic Role of TAZ in Human Non-Small Cell Lung Cancer. Clinical Cancer Research, 20, 4660–4672. doi:10.1158/1078-0432.CCR-13-3328

Pearce, R. G., Setzer, R. W., Strope, C. L., Sipes, N. S., & Wambaugh, J. F. (2017). httk : R Package for High-Throughput Toxicokinetics. Journal of Statistical Software, 79(4). doi:10.18637/jss.v079.i04

Peck, D., Crawford, E. D., Ross, K. N., Stegmaier, K., Golub, T. R., & Lamb, J. (2006). A method for high-throughput gene expression signature analysis. Genome Biology, 7(7). doi:10.1186/gb-2006-7-7-r61

Pommier, Y. (2013). Drugging topoisomerases: Lessons and Challenges. ACS Chemical Biology. doi:10.1021/cb300648v

Pommier, Y., Barcelo, J. M., Rao, V. A., Sordet, O., Jobson, A. G., Thibaut, L., … Redon, C. (2006). Repair of Topoisomerase I-Mediated DNA Damage. Progress in Nucleic Acid Research and Molecular Biology. doi:10.1016/S0079-6603(06)81005-6

Richard, A. M., Judson, R. S., Houck, K. A., Grulke, C. M., Volarath, P., Thillainadarajah, I., … Thomas, R. S. (2016). ToxCast Chemical Landscape: Paving the Road to 21st Century Toxicology. Chemical Research in Toxicology. doi:10.1021/acs.chemrestox.6b00135

Ritchie, M. E., Phipson, B., Wu, D., Hu, Y., Law, C. W., Shi, W., & Smyth, G. K. (2015). limma powers differential expression analyses for RNA-sequencing and microarray studies. Nucleic Acids Research, 43(7), e47. doi:10.1093/nar/gkv007

Schmidt, C. W. (2009). Tox21: New dimensions of toxicity testing. Environmental Health Perspectives, 117(s8). doi:10.1289/ehp.1181c4

Shimizu, H., Yisireyili, M., Higashiyama, Y., Nishijima, F., & Niwa, T. (2013). Indoxyl sulfate upregulates renal expression of ICAM-1 via production of ROS and activation of NF-??B and p53 in proximal tubular cells. Life Sciences, 92(2), 143–148. doi:10.1016/j.lfs.2012.11.012

Smyth, G. K. (2005). Limma: linear models fro microarray data. In Bioinformatics and Computational Biology Solutions using R and Bioconductor (pp. 397–420). doi:10.1007/0-387-29362-0_23

Soldatow, V. V. Y., Lecluyse, E. E. L., Griffith, L. L. G., & Rusyn, I. (2013). In vitro models for liver toxicity testing. Toxicology Research, 2(1), 23–39. doi:10.1039/C2TX20051A.In

Sonja, H., Castelo, R., & Guinney, J. (2014). GSVA : The Gene Set Variation Analysis package for microarray and RNA-seq data. Bioconductor.org, 1–20. Retrieved from http://www.bioconductor.org/packages/release/bioc/vignettes/GSVA/inst/doc/GSVA.pdf

Stanford, E. A., Ramirez-Cardenas, A., Wang, Z., Novikov, O., Alamoud, K., Koutrakis, P., … Sherr, D. H. (2016). Role for the Aryl Hydrocarbon Receptor and Diverse Ligands in Oral Squamous Cell Carcinoma Migration and Tumorigenesis. Molecular Cancer Research, 14(8), 696–706. doi:10.1158/1541-7786.MCR-16-0069

Subramanian, A., Narayan, R., Corsello, S. M., Peck, D. D., Natoli, T. E., Lu, X., … Golub, T. R. (2017). A Next Generation Connectivity Map: L1000 Platform and the First 1,000,000 Profiles. Cell, 171(6), 1437–1452.e17. doi:10.1016/j.cell.2017.10.049

Subramanian, A., Tamayo, P., Mootha, V. K., Mukherjee, S., Ebert, B. L., Gillette, M. A., … Mesirov, J. P. (2005). Gene set enrichment analysis: A knowledge-based approach for interpreting genome-wide expression profiles. Proceedings of the National Academy of Sciences, 102(43), 15545–15550. doi:10.1073/pnas.0506580102

Tice, R. R., Austin, C. P., Kavlock, R. J., & Bucher, J. R. (2013). Improving the human hazard characterization of chemicals: A Tox21 update. Environmental Health Perspectives. doi:10.1289/ehp.1205784

Tomasetti, C., Li, L., & Vogelstein, B. (2017). Stem cell divisions, somatic mutations, cancer etiology, and cancer prevention. Science, 355(6331), 1330–1334. doi:10.1126/science.aaf9011

Tomasetti, C., & Vogelstein, B. (2015). Variation in cancer risk among tissues can be explained by the number of stem cell divisions. Science, 347(6217), 78–81. doi:10.1126/science.1260825

Uehara, T., Minowa, Y., Morikawa, Y., Kondo, C., Maruyama, T., Kato, I., … Urushidani, T. (2011). Prediction model of potential hepatocarcinogenicity of rat hepatocarcinogens using a large-scale toxicogenomics database. Toxicology and Applied Pharmacology, 255(3), 297–306. doi:10.1016/j.taap.2011.07.001

Wang, L., & Eastmond, D. A. (2002). Catalytic inhibitors of topoisomerase II are DNA-damaging agents: Induction of chromosomal damage by merbarone and ICRF-187. In Environmental and Molecular Mutagenesis (Vol. 39, pp. 348–356). doi:10.1002/em.10072

Wang, Y., Fan, Y., & Puga, A. (2010). Dioxin exposure disrupts the differentiation of mouse embryonic stem cells into cardiomyocytes. Toxicological Sciences : An Official Journal of the Society of Toxicology, 115(1), 225–37. doi:10.1093/toxsci/kfq038

Zhang, L., & Gong, F. (2016). Involvement of USP24 in the DNA damage response. Molecular & Cellular Oncology, 3(1), e1011888. doi:10.1080/23723556.2015.1011888

